# Modelling liver cancer microenvironment: novel 3D culture system as a potential anti-cancer drug screening tool

**DOI:** 10.1101/2021.11.04.467266

**Authors:** Ala’a Al Hrout, Karla Cervantes-Gracia, Richard Chahwan, Amr Amin

## Abstract

The tumor microenvironment and its contribution to tumorigenesis has been a focal highlight in recent years. A two-way communication between the tumor and the surrounding microenvironment sustains and contributes to the growth and metastasis of tumors. Progression and metastasis of hepatocellular carcinoma have been reported to be exceedingly influenced by diverse microenvironmental cues. In this study, we present a 3D-culture model of liver cancer to better mimic *in vivo* tumor settings. By creating novel 3D co-culture model that combines free-floating and scaffold based 3D-culture techniques of liver cancer cells and fibroblasts, we aimed to establish a simple albeit reproducible *ex vivo* cancer microenvironment model that captures tumor-stroma interactions. The model presented herein exhibited unique gene expression and protein expression profiles when compared to 2D and 3D mono-cultures of liver cancer cells. Our results showed that in vivo like conditions cannot be mimicked by simply growing cancer cells as spheroids, but by co-culturing them with 3D fibroblast with which they were able to cross-talk. This was evident by the upregulation of several pathways involved in HCC, and the increase in secreted factors by co-cultured cancer cells, many of which are also involved in tumor-stroma interactions. Compared to the conventional 2D culture, the proposed model exhibits an increase in the expression of genes associated with development, progression, and poor prognosis of HCC. Our results correlated with an aggressive outcome that better mirrors *in vivo* HCC, and therefore, a more reliable platform for molecular understanding of HCC and possibly better anti-cancer drug screening.

## INTRODUCTION

Cancer is a multi-factorial disease, arising from normal cells, primarily through abnormal cellular proliferation and progressive mutation load. Tumor cells, however, represent only one aspect of tumorigenesis. The tumor milieu is composed of a dynamic network of non-malignant cellular components, non-cellular components, signaling molecules, and extracellular matrix (ECM) [1,2], which collectively forms the tumor microenvironment (TME). A dynamic two-way communication between the tumor and the surrounding milieu, sustains and contributes to tumor growth and metastasis [3]; thereby highlighting the key role the TME plays in tumor progression [1,4]. In addition, many studies have reported the positive role of the TME in restraining tumor initiation and progression at initial stages of carcinogenesis [5], and how “reprogramming” the TME in the later stages holds a great potential for developing effective cancer treatments [1].

Fibroblasts are generally considered the predominant cellular TME component. Whilst normally in an “inactive” quiescent state; fibroblasts recruited to the tumor site are constantly activated by the tumor through paracrine signaling, after which they are transformed into cancer-associated fibroblasts (CAFs) [6]. Once the CAF transition is triggered, paracrine signaling is no longer needed [7]. These transformed CAFs become distinct in their morphology and function from normal fibroblasts [7], most likely due to their rewiring by tumor signaling. CAFs possess higher ability to proliferate [8], be tumor proximal, and evade apoptosis [9]. But the molecular mechanisms mediating this process remains elusive. CAFs contribute significantly to tumorigenesis; partly through suppressing immune responses, secreting growth factors, cytokines, and proangiogenic factors [10]. In addition, CAFs contribute to tumorigenesis through secreting ECM proteins and degrading matrix metalloproteinase (MMPs), which together, give CAFs their ECM remodeling ability [10]. CAFs, therefore, have potential as therapeutic targets [11].

A tumor has an increasing demand for oxygen and nutrients to support its progression. When the demand for oxygen remains unmet, low oxygen hypoxic conditions ensue [12]. To survive, tumor cells activate the hypoxia-inducible factor 1 (HIF1) [12], which in turn activates the transcription of a group of genes through binding to their hypoxia-response elements to promote the survival of tumor cells [13]. HIF-1 targeted genes significantly contribute to tumor angiogenesis, metastasis, adhesion, metabolism, and pH regulation [13]. Moreover, many studies have highlighted the role of hypoxia in recruiting stromal components to the TME [14], ECM composition, and metastatic remodeling [15] Hepatocellular carcinoma (HCC), is the fifth most common cancer and is the fourth cause of cancer-related death worldwide [16]. HCC has a very poor prognosis with only five-year survival rate [17]. HCC progression is influenced by the liver microenvironment such as altered stromal cells [18]. These cells deposit ECM proteins causing fibrosis that then progresses to cirrhosis with a prevalence of 80-90% [18]; suggesting a crucial role of ECM build-up in HCC progression [19]. Hypoxia represents a driving force for HCC progression, and is associated with poor prognosis [20]. HIF-mediated gene expression contributes to different aspects of HCC metastasis, such as epithelial mesenchymal transition (EMT) [21], invasion of the ECM, and metastasis [22]. Yet the molecular mechanisms governing stromal and tumor cell interactions within the TME of HCC under hypoxic conditions [14] remains unclear.

To reflect the complexity and dynamic nature of tumor cell biology, a physiologically relevant model is needed. Especially, when it comes to drug discovery and identifying effective therapeutic targets. To simulate *in vivo* environment, *in vitro* two-dimensional (2D) cell culture is typically assembled by growing cells *commonly* on a plastic substrate in an adherent monolayer. However, distortion of spatial arrangement of cells in 2D culture changes cell-cell and cell-matrix interactions [23], and most importantly, alters the response of cells to certain drugs and treatments [24]. That is why, 3D cultured cells better recapitulates *in vivo* architecture of tumors and exhibits gene expression closer to that of *in vivo* tumors [23]. One very common 3D cultured cells model is the spheroid; a micro cell cluster sphere [25]. The nature of this three-dimensional multicellular model is what makes it an attractive tool to simulate solid tumors *in vitro* as it is composed of three regions, a highly proliferative outer region, a middle quiescent region, and a hypoxic core region [26]. Such compartmentalization creates diffusional gradients of oxygen, nutrients, and tested drugs among all three regions of the spheroid, which is also characteristic of solid tumors [27].

We aimed at modeling the basic TME of liver cancer by mimicking certain aspects of in vivo tumors, such as three-dimensionality of tissue, hypoxia, and heterogeneity of tumors. Five groups were designed to reflect each element, group 1 is a control group for comparison purposes which consists of 2D mono-cultures of liver cancer cells. Group 2 is also 2D mono-cultures of liver cancer cells but induced for hypoxia chemically. Group 3 on the other hand is like group 2 but additionally includes conditioned media from 2D fibroblasts to reflect a oneway co-culture system. Group 4 and 5 are 3D cultures of liver cancer cells that exhibit hypoxia physiologically due to culturing conditions. However, group 5, which is our proposed model, includes a 3D culture of fibroblast in a separate insert, reflecting a two-way co-culture system. Our findings reported herein demonstrate that our proposed model of group 5 reflects many aspects of in vivo settings and signaling pathways, promoting it as a potential platform for further studies of drug efficacy in vitro and understanding the communication between cancer and the stroma in liver cancer.

## MATERIALS AND METHODS

### Co-culture Systems

For the 2D co-culture, a one-way communication system was followed. Briefly, HepG2 and SV-80 cells were seeded at a density of 1× 10^6^ in conventional 2D culturing flasks in DMEM media. Flasks were incubated at 37°C in 5% CO_2_ humidified incubator for about 24 hours. Conditioned media of SV-80 cell line was collected, centrifuged to collect any cellular debris, and applied to HepG2 cells, which were incubated with the conditioned media for 48 hours.

For the 3D co-culture, a two-way communication system was followed. Briefly, HepG2 and SV-80 3D cultures were prepared separately. Prior to co-culture, 6-well plates were coated with 1.5% agarose, and allowed to set and cool before transferring HepG2 spheres to the bottom of the coated plates, and the inserts containing SV-80 3D culture were placed on top. HepG2 and SV-80 3D cultures were incubated for 48 hours.

The remaining materials and methods are included in supplementary data.

## RESULTS

### Comparing 2D, mono-3D, and co-3D cultures and their gene expression

Fibroblasts grown as a 2D culture exhibit an elongated morphology. However, growing fibroblasts as 3D culture in porous scaffolds alters their morphology to be more rounded. HepG2 cells grown in 3D were monitored over a period of 5 days to assess formation of tight spheroids with a smooth surface (Fig. 1A-B. HepG2 2D cultures were treated with increasing concentrations of CoCl_2_ (100-400 μM) to assess cellular viability under hypoxia-mimicking conditions (Fig. 1C). Treatment of HepG2 with CoCl_2_ did not affect cellular viability significantly at doses of 100 and 200 μM of CoCl_2_. However, a highly significant difference (*p* < 0.001) was noted at a dose of 400 μM CoCl_2_. To confirm the induction of hypoxia in CoCl_2_-treated HepG2 2D cultures, protein expression of HIF1α was assessed using western blot. Treatment with 200 μM of CoCl_2_ for 6 hours did not induce HIF1α expression. However, by increasing the dose to 300 μM, HIF1α expression was detected in HepG2 cells (Fig. 1D). Immunofluorescence was then used to detect the cellular localization of HIF1α, showing that CoCl_2_ also affects HIF1α translocation to the nucleus (Fig. 1E), where it can bind to hypoxia-response elements (HREs).

**Figure 1.**
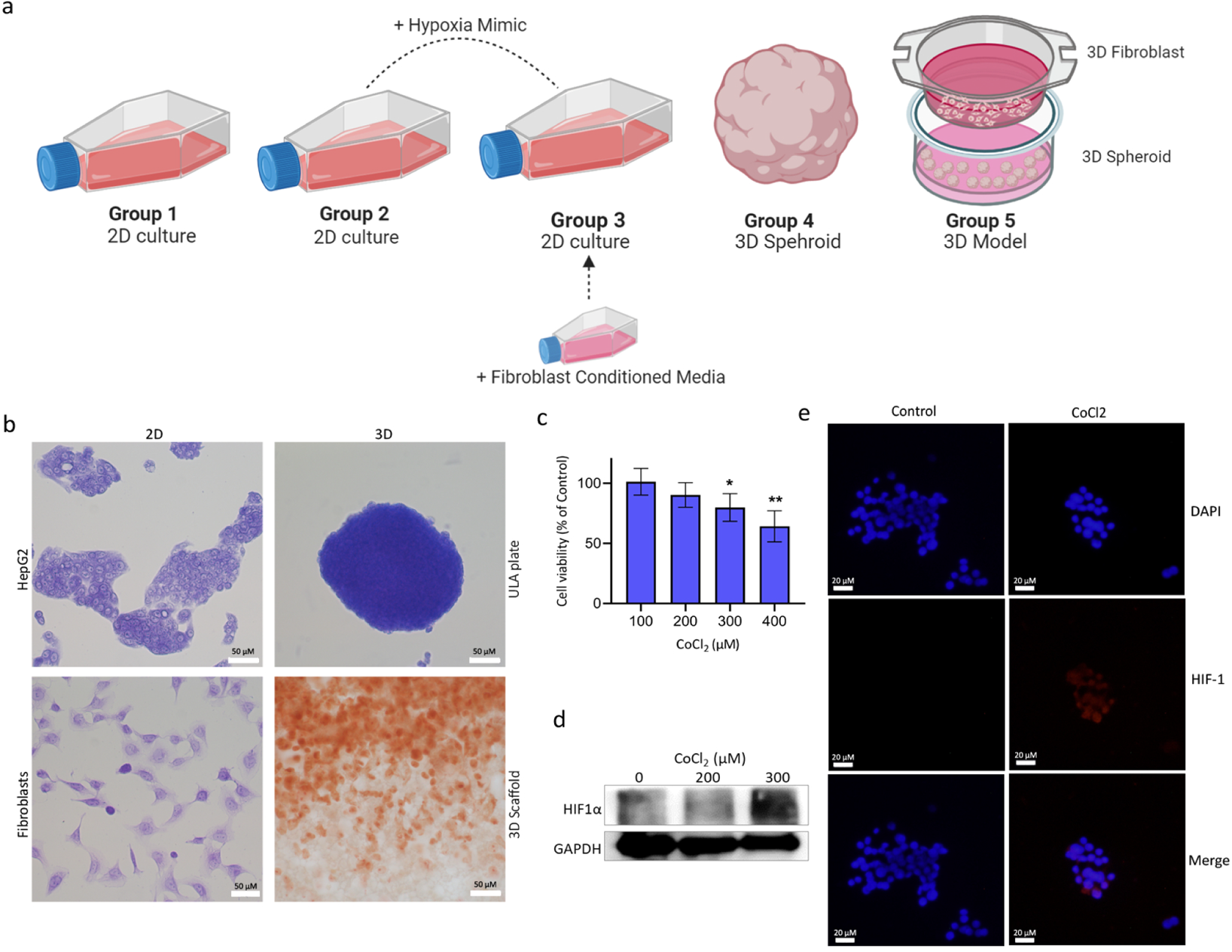
Experimental model and initial validations. a) Schematic depiction of experimental groups. b) Bright field images of 2D/ 3D HepG2 and fibroblasts cultures fixed and stained with crystal violet and 3D fibroblasts fixed and stained with neutral red (scale bar = 50 μm). c) Cell viability of HepG2 cells after treatment with increasing concentrations of CoCl_2_ for 6 hours (p < 0.01). d) HIF1-α protein expression was detected in CoCl_2_ HepG2-treated cells after 6 hours of incubation. Uncropped blots are shown in Fig. S1. e) Immunofluorescence detection of HIF1-α expression and localization in HepG2 cells incubated with or without CoCl_2_ (scale bar = 20 μm).

### Differential gene expression analysis

To better assess in vivo mimicking capabilities of our 5 groups, RNA from the denoted samples (Fig. 1A) were extracted and then sequenced via the Illumina platform. Significant differentially expressed genes (DEGs) were determined at a cutoff of Log2FC of 1 and FDR < 0.05. A hierarchical clustering Heatmap was used to explore changes in global gene expression across our 5 models (Fig. 2 A). Interestingly, group 5 clustered separately from the rest, revealing a fundamental and distinctive change in gene expression. Principal component analysis (PCA) based on the global gene expression showed a similar outcome, where group 5 segregated at the opposite extreme of all other groups, with the conventional HepG2 culture (group 1) being the most distant (Fig. 2B), suggesting fundamental differences between 2D, monocellular 3D, and multicellular 3D models.

**Figure 2.**
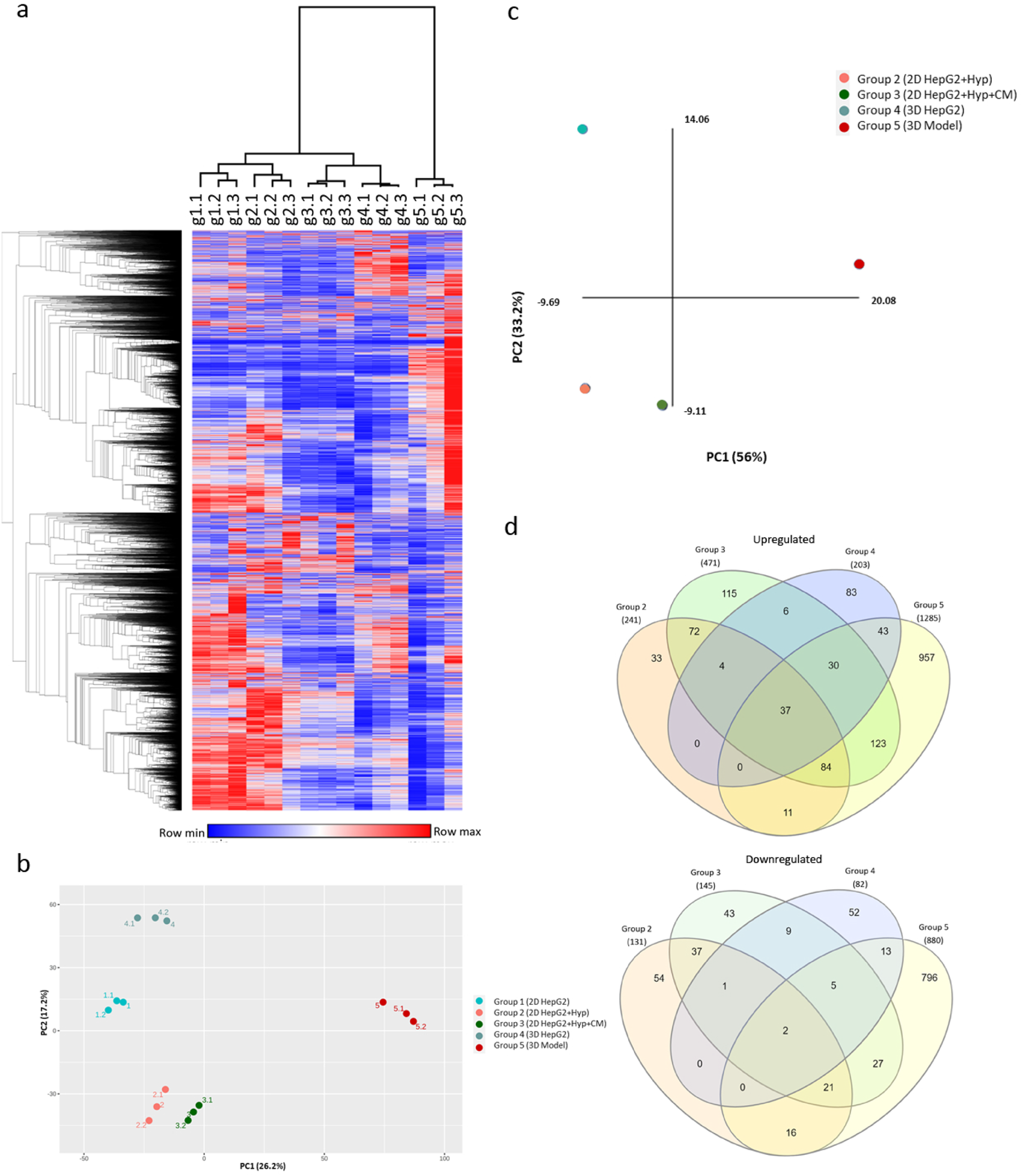
Computational analysis of the RNA-Seq of our group cohorts. a) Hierarchical clustering heatmap of global expression in all groups (in triplicates) was generated with Morpheus using default settings. B) PCA plot based on global expression in all groups (in triplicates) was generated using NetworkAnalyst c) PCA plot based on significant DEGs of group 2,3,4,5 in comparison to group 1. d) Venn diagrams showing the common and unique upregulated genes and downregulated genes of groups 2, 3, 4, and 5. The interactive diagrams can be accessed online using the InteractiVenn (http://www.interactivenn.net) and uploading supplementary files 1 & 2c)

Significant differentially expressed genes (DEG) were determined relative to group 1 using RNA-seq 2G. Group2 resulted in significant upregulation of 243 genes and downregulation of 131 genes. Group3 increased the number of significantly DEGs to 474 upregulated and 145 downregulated genes. When comparing gene expression profile of group 4 to group 1, 203 genes were significantly upregulated, whereas 82 genes were downregulated. Group 5 dramatically changed the gene expression of HepG2 by significantly upregulating 1291 and downregulating 880 genes in group 5 in comparison to group 1. PCA plot based on the DEGs showed similar results to the global expression PCA plot, whereby group 5 was segregating separately from the remaining groups (Fig. 2C). In Addition, significant DEGs signatures, unique to each group, were identified (Fig. 2D). Group 5 exhibited the highest number in unique significant DEGs among all other groups, at 957 upregulated genes and 796 downregulated genes. A full List of common and unique significantly DEGs of each group is available in supplementary files 1 and 2.

Canonical pathways associated with significant DEGs in different culturing conditions were analyzed using gene-based enrichment analysis by XGR. As expected, culturing HepG2 cells under hypoxia-mimicking conditions (group 2 and 3) upregulated genes involved in hypoxia-inducible factor-1 alpha (HIF1-α) and hypoxia-inducible factor-2 alpha (HIF2-α) pathways, and networks downstream of these pathways. This was even observed in 3D spheroid cultures (group 4 and 5) despite not being treated with a hypoxia inducing agent, indicating the formation of hypoxic core in 3D culture spheroids. Culturing HepG2 cells with only fibroblasts conditioned media (group 3) or alternatively with 3D culture of fibroblasts (group 5) upregulated genes involved in integrin family cell surface interactions, interleukin-6 (IL6) mediated signaling events (Table S1).

### In-depth pathway functional analysis reveals pathways associated with HCC progression

To further understand the role of significant DEGs in each group, we applied clustering analysis using ClueGO/CluePedia as described previously [28]. By combining Gene Ontology (GO) terms, KEGG and Wiki pathways, ClueGo/CluePedia create a better interpretation of the pathways associated with the list of input genes [29]. Hypoxia mimicking conditions in group 2 induced cellular responses to hypoxia and HIF-1 signaling pathway, as well as other signaling pathways reported to promote HCC including NRF2, FOXO, and p53 pathways [30,31] (Fig. 3A-B). Similarly, inducing hypoxia in group 3 resulted in inducing HIF-1 signaling pathways. However, with the addition of fibroblast conditioned media, group 3 DEGs-associated processes were enriched in 3 out of 5 pathways previously reported in KEGG analysis of HCC patients’ tissues including i) complement and coagulation cascades, ii) focal adhesion, and iii) ECM-receptor interaction [32] (Fig. 3A). DEGs-associated processes of HepG2 3D culture alone (group 4) resembled some of group 2 such as hypoxia and NRF2 signaling pathways, but also some of group 3 such as G3 such as complement and coagulation cascades (Fig. 3A). Additionally, DEGs-associated processes of group 4 included steroid hormone biosynthesis process and estrogen signaling pathway, which have been linked to HCC progression [33] (Fig. 3A). Group 5 shared some pathways with the other groups such as hypoxia, focal adhesion, glycolysis/ gluconeogenesis, and estrogen signaling pathway. In addition, DEGs-associated processes of group 5 were significantly enriched in HCC-promoting pathways including oncostatin M signaling pathway, insulin signaling pathway and aryl hydrocarbon receptor pathways [34,35] (Fig. 3B).

**Figure 3.**
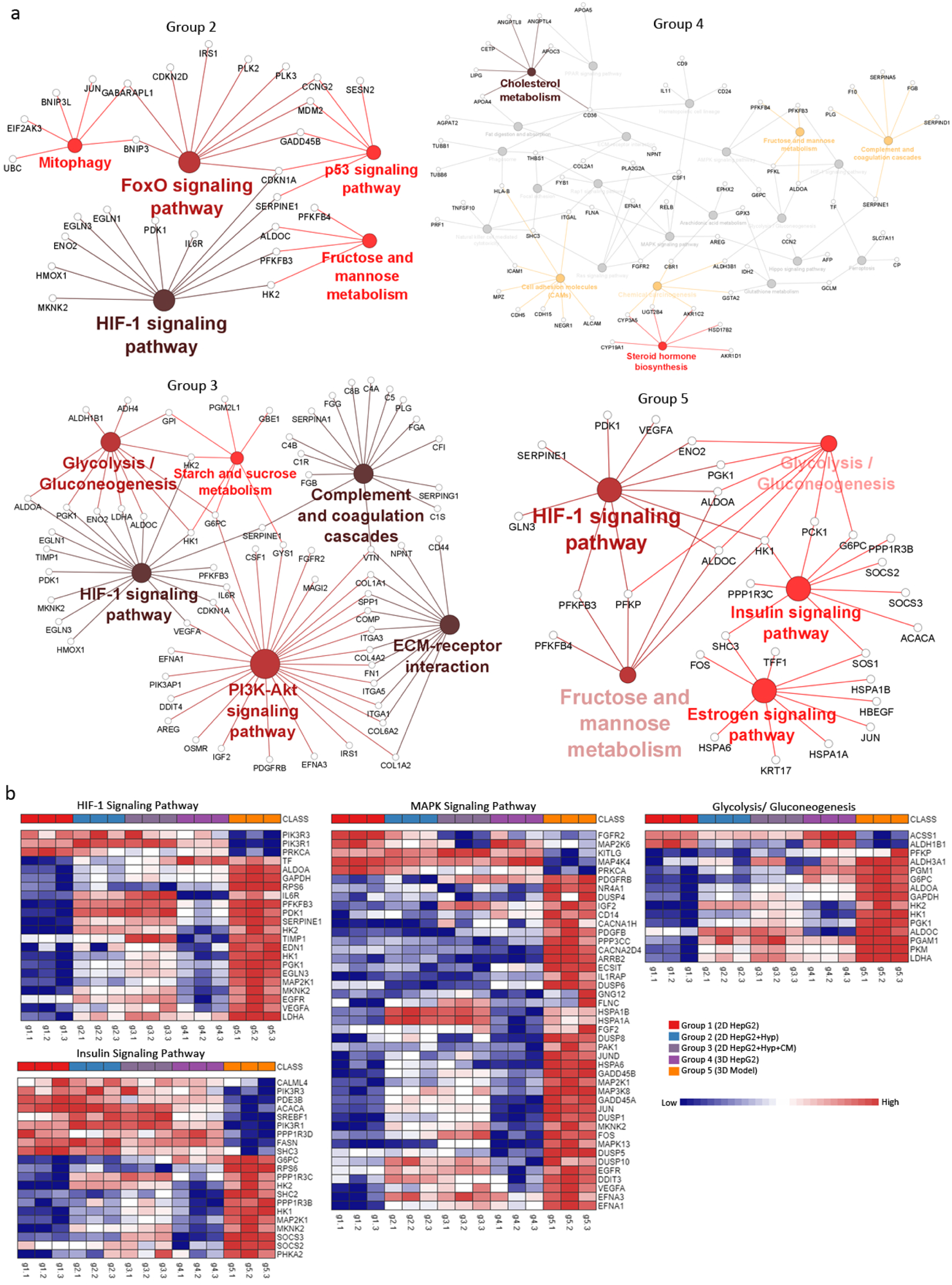
Pathway clustering analysis and heatmap representation of experimental groups. a) Pathway term clustering based on KEGG pathway maps based on DEGs of experimental groups. All groups are compared to G1 as the control condition. Node size of pathway terms resemble the number of associated genes to it. The stronger the node color the more significant a cluster is. b) ORA heatmaps of enriched genes in denoted pathways based on KEGG from all groups. Heatmaps were generated using NetworkAnalyst.

After identifying the main pathways involved in group 5 through ClueGO/CluePedia clustering, Insulin signaling pathway was further analyzed and visualized in PathVisio due to its documented relevance in HCC [36] and to gain insights of significant physiological changes occurring within a specific pathway. Genes below FDR 0.05 (5871 hits) were imported into PathVisio [37] to identify trends in regulation within this particular pathway and other chained events embedded within associated pathways. Inconsistencies within these maps were excluded from the final pathway (Fig. 4). The Insulin signaling pathway in group 5 was found to mainly lead to activation of genes involved MAPK signaling and this trend converges with hypoxia signaling input, both leading to the production of VEGFB, VEGFA, PGF, growth factors known to be involved in angiogenesis and tumor invasion [38]. Tumor invasion can also be promoted via Ras and its downstream pathways of MAPK and RALB, which were all shown to be upregulated in group 5. Some of the genes involved in the pathway shown previously have been validated with qPCR (Fig. 5A).

**Figure 4.**
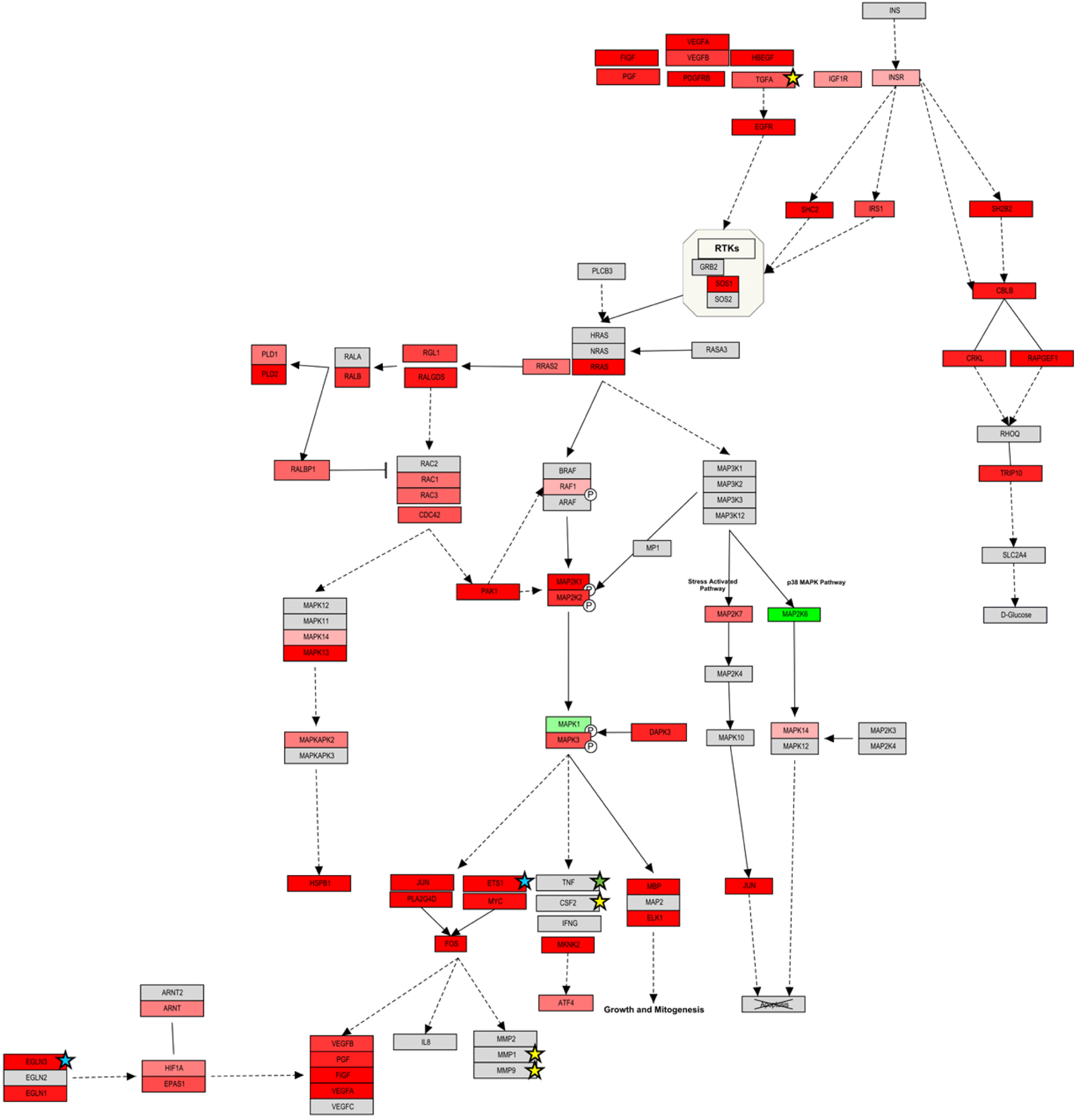
Customized Insulin signaling pathway map of Group 5 DEGs. Insulin signaling pathway identified as significant through ClueGO/CluePedia pathway term clustering analyzed and customized through Pathvisio. Red boxes: up regulated genes; green boxes: down regulated genes; gray boxes: genes not found within the DEGs. Color intensity resembles the deregulated status of the gene from the LogFC scale (between a 1 to −1 LogFC). A clear up-regulated trend was identified that resembles the activation of the pathway and the main genes involved within this mechanism. This customized pathway map delineates the line of significance from G5 Insulin signaling pathway. These results back-up and *validate ClueGO/CluePedia analysis. Yellow and blue stars highlight the genes/proteins that were further validated either through antibody arrays or qPCR respectively*.

**Figure 5.**
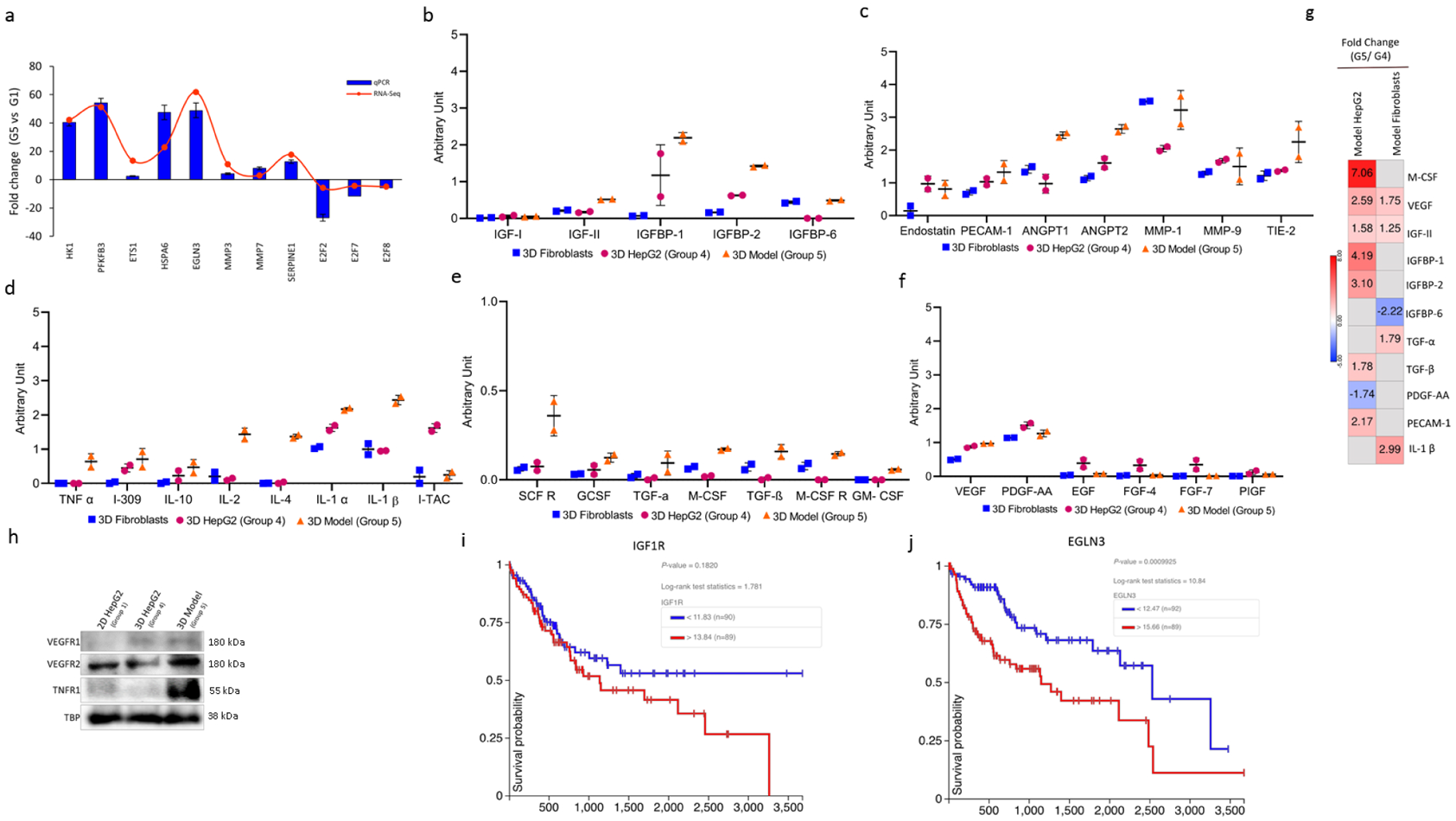
Confirmation studies using genomic, proteomic, and clinical analyses. a) Validation by qPCR. RNA-Seq based expression was plotted against qPCR-based expression. qPCR data are represented as means of fold change ± SD. b-f) levels of different secreted factors in 3D mono- and co-cultures analyzed by antibody arrays. Experimental scheme is outlined in Fig. S2 g) fold change of expression of HepG2 and Fibroblasts before and after co-culture for markers shown in panels b-f based on RNA-seq data. h) western blotting of total cell lysates of 2D or 3D HepG2 cultures. Uncropped blots are shown in Fig. S3. i-j) Kaplan–Meier curves of progression-free survival of IGF1R and EGLN3 in HCC patients.

### Co-culture of Fibroblasts and HepG2 3D cultures enriches the secretome in pro-tumor factors

To identify signaling factors playing a role in the crosstalk between cancer cells and fibroblasts, the secretome of group 4 and 5 was analyzed for factors involved in angiogenesis, invasion, and metastasis, in addition to other factors in the insulin signaling pathway, secretome of 3D fibroblasts was included as a control (Fig S2, Fig. 5B-F). Most factors were dramatically increased in the secretome of group 5 (from both fibroblasts and HepG2 cells) when compared to 3D mono-cultures. Co-culturing HepG2 spheroids with 3D culture of fibroblasts increased the levels of insulin signaling pathway factors like IGF-II, IGFBP1, IGFBP2 (Fig 5B). In addition, levels of different angiogenesis and cytokines that are involved in the cross-talk in the TME were highly increased in the setting of group 5 (Fig 5C & F).

As the secretome collected from group 5 is shared among fibroblasts and HepG2 cells, we wanted to confirm the source of the secreted factors. To that end we compared gene expression of 3D HepG2 or 3D fibroblasts before and after co-culture (group 5 over group 4) (Fig. 5G). Differential gene expression analysis revealed that M-CSF, IGFBP1, IGFBP2, TGF-β, and PECAM-1 were upregulated in HepG2 cells after co-culture, while they were not differentially expressed in fibroblasts. On the other hand, TGF-α, and IL-1β were only differentially expressed in fibroblasts after co-culture. VEGF and IGF-II were differentially expressed in both HepG2 cells and fibroblasts, being more upregulated in HepG2 cells (Fig. 5G). Despite that VEGF is upregulated more than 2.5 folds on the RNA level in HepG2 cells after co-culture (Fig. 5G), there was no noticeable difference in the secreted VEGF from 3D HepG2 cells before and after co-culture (Fig. 5F). Nonetheless, VEGFR2 was more expressed in group 5 in comparison to group 1 and group 4; while VEGFR1 was higher in the 3D cultures, but no noticeable difference between the two (Fig. 5H). TNFR1 was also dramatically higher in group 5 when compared to both group 1 and group 4 (Fig. 5H). Kaplan-Meier curves of genes within our main pathway of interest were analyzed. IGF1R and EGLN3 showed a significant correlation between expression and prognosis (Fig. 5 I-J). Both genes were shown to be involved in insulin signaling pathway (Fig. 4).

### Pathway complementation analysis highlights regulatory miRNAs and prognosis markers

To better understand the changes elicited by the 3D co-culture conditions on HepG2 cells, we investigated the effect of miRNA regulation given their role in affecting *de novo* or modulating established gene expression in tumors [39,40]. Based on miRNA-gene interaction analysis of significant DEGs in group 5, miR-335 was the top regulatory miRNA with the highest number of connections to other genes in the network (Fig. S4 A). Other miRNAs were predicted, top 10 are shown in Fig. S4 B with literature citations (full list is available in Table S2).

## DISCUSSION

### Establishment of 3D culture models

We investigated the bilateral effect of the cellular architecture and milieu on the properties and phenotypic responses of the whole cellular microenvironment of tumor cells. Liver cancer microenvironment was modeled using 2D and 3D mono- and co-culture systems. The transcriptome of five models was profiled to determine the effects of each condition. Each one of the culturing conditions mimicked an aspect of the TME. Based on our clustering methods, group 5 clustered away all the other groups, specifically group 1, in both the PCA plot and hierarchical cluster heatmap (Fig. 2 A-C). This is in line with the Venn diagram, where group 5 had the highest number of unique genes that are not shared with other groups (Fig. 2D). This clear distinction suggests a fundamental difference in gene expression between the proposed model of group 5 and all other culturing conditions, signifying a considerable change in gene expression as a result of 3D co-culture of cancer spheroids with fibroblasts.

### Mimicking hypoxic conditions is not sufficient to mimic in vivo conditions

Treatment with hypoxia mimicking agent significantly upregulated genes associated with p53 and AP-1 networks (Table S1). AP-1 proteins (*i*.*e*. Jun, Fos, and ATF families) are activated by hypoxia and are frequently deregulated in cancer [41]. Deregulation of ATF-2 and its network have been implicated in liver development, regeneration, and cirrhosis [42]. In addition, ATF-2 has been reported to play a role in HCC resistance to sorafenib, *in vivo* [43]. The presented hypoxia mimicking conditions also induced several metabolism pathways including fructose, mannose, starch, and sucrose metabolism; as well as glycolysis and gluconeogenesis (Fig. 3). Nonetheless, without treatment with CoCl_2_, HepG2 spheroids (group 4 & 5) upregulated genes were associated with HIF1-α and HIF2-α (Fig. 3A; Table S1). This is consistent with studies promoting cancer spheroids as candidate solid tumor model, as they recapitulate many aspects of *in vivo* tumors including hypoxia [44]. Hypoxia-mediated pathways in turn promote survival, angiogenesis, invasion, and metastasis [45]. Hypoxia also plays a role in lipid and steroid metabolism [46]. Steroids promote tumor immune evasion by suppressing T cell activation and subsequently effecting immune-therapy outcome [47]. This is consistent with induction of steroid synthesis and estrogen signaling pathways in group 4 and 5 (physiological hypoxia), but not group 2 and 3, suggesting that chemically mimicking hypoxia is not sufficient to recapitulate a more encompassing hypoxic condition (Fig. 3; Table S1). Taken together, these results demonstrate how adjusting HepG2 cells from 2D to 3D culture introduces hypoxia and its associated hallmarks, which better represents *in vivo* cancer conditions.

### Reconstitution of the dynamic bilateral interaction between cancer and stromal cells through 3D co-culture system

Studying the interaction between cancer cells, stromal fibroblasts, and their contribution to tumorigenesis remains a challenge. We, therefore, created a simplified system to mimic stroma-tumor interaction. HepG2 cells were either cultured with fibroblasts conditioned media only (group 3) to mimic one-way interaction or cultured with fibroblasts in a 3D-culture system (group 5) to mimic a two-way communication. When cultured with fibroblasts, HepG2 upregulated genes were associated with integrin cell surface interactions, in addition to urokinase-type plasminogen activator (uPA) and uPAR-mediated signaling. Integrin signaling pathway (ISP) regulates the interaction with the extracellular environment in response to intracellular cues [48] β1 integrins are overexpressed in many tumors, and blocking their signaling transduction reduces survival and tumorgenicity of many cancers, in 2D and 3D *in vitro* cultures, and *in vivo* [49–52]. ISP transduction has been reported to be promoted by other cell surface proteins, such as urokinase receptor (uPAR) [53]. Suppressing uPAR expression or disruption uPA/uPAR interaction have been reported to inhibit tumor progression and metastasis [54]. In addition, canonical pathways associated with the upregulated genes in group 3 and 5 included IL-6 mediated signaling. This is consistent with Integrin increased signaling, where enhanced IL-6/STAT3 signaling is promoted by β1-Integrin pathway [55,56]. IL-6 overexpression has been reported in many cancers, including HCC, where it is suggested to promote the transition of fibroblasts to CAFs [57]. Indeed, studies have shown CAFs as the main source of IL-6, promoting survival, migration, invasion, angiogenesis and stemness in colorectal, gastric, and liver cancer cells [58]. By co-culturing HepG2 cells with fibroblasts, we were able to recapitulate signaling pathways essential in tumor-stroma crosstalk, creating a more reliable model to study complex TME interactions.

### Recapitulating signaling pathways in HCC through 3D co-culture system

Insulin/IGF signaling pathway is known to be activated in many cancers including HCC. Studies have shown its essential role in carcinogenesis and metastasis [59]. The insulin pathway was generally activated in 3D model (group 5) in comparison to other culturing methods. Many of the factors involved in insulin pathway – in addition to factors involved in hypoxia, angiogenesis, and MAPK signaling – were upregulated in our study (Fig. 3 & 4); which was confirmed on the transcriptome and secretome levels (Fig. 5). HK1 is dramatically upregulated in group 5 (Fig. 5a). HK1 shows involvement in glycolysis, HIF, and insulin signaling pathways (Fig. 3A-B), a role that has been reported in various studies; where HK1 contributes to glycolysis, proliferation, migration, and invasion of HCC [40,60,61]. Similarly, PFKFB3, a direct target of HIF1, is upregulated in all groups in comparison to group 1 and regulates glucose metabolism and promotes cancer progression and growth [62]. Overexpression of PFKFB3 is associated with poor prognosis of HCC, and its inhibition resulted in suppression of HCC growth *in vitro* and *in vivo* [63] and reversed the in vitro sorafenib-resistance of HCC cells [64]. EGLN3, that mediates crosstalk between hypoxia and insulin signaling pathways [65,66], was upregulated in HCC hypoxic settings [67,68]. Consistently, EGLN3 was a significant factor driving hypoxia and insulin pathways in our proposed model (group 5). Culturing HepG2 under 3D co-culture conditions also enhanced the expression of genes promoting angiogenesis, migration, and invasion such as SERPINE1, ETS1, MMP3, and MMP7. ETS1, MMP3, and MMP7 are only upregulated in group 5 in comparison to group 1. ETS1 is involved in upregulating hypoxia-target genes such as MMP3, and MMP7 [69,70]. Downregulation of ETS1 was reported to inhibit metastasis and invasion of liver cancer cell lines [71]. Similarly, Expression of MMP3 and MMP7 is correlated with enhanced metastatic phenotype, where their inhibition suppressed invasion and migration of HCC cells [18]. SERPINE1 was distinctively upregulated in all groups compared to group 1, yet, with the highest fold change exhibited in group 5. Increased expression of PAI-1 (encoded by SERPINE1) is correlated with aggressive cancers and poor prognosis, where it is also associated with migration, invasion, and angiogenesis in HCC tissue [72]. Culturing HepG2 under 3D co-culture conditions also downregulated the expression of genes involved in cell cycle regulation and survival, including E2F2, E2F7, and E2F8. E2F transcription factors were only differentially expressed in group 5, where they were found to be downregulated in comparison to group 1, as confirmed by qPCR (Fig. 5a). Downregulation of E2F2, E2F7, and E2F8 prevents cell cycle arrest and enhances clonogenic survival [73]. Taken together, these findings indicate that culturing HepG2 in a 3D co-culture system (group 5) results in enhanced migration, invasion, metastasis, and angiogenesis, and successfully recapitulating many signaling pathways that are important in HCC *in vivo*.

### Pathway complementation analysis highlights the involvement of regulatory miRNAs

The cross-talk between cancer cells and other cells in the TME is also partly mediated through expression of miRNA and/or release of miRNAs through extracellular vesicles (EVs) [74]. Gene-miRNA interaction network analysis of significant DEGs of group 5 revealed a number of predicted regulatory miRNAs (Fig. S4, full list in Table S2), many of which are signature of HCC deregulated miRNAs [39,60]. For instance, miR-335 had the highest number of connections in group 5 (Fig. S4 a). Serum of HCC patients undergoing TACE was analyzed for circulating miRNAs level, where miR-335 level was associated with significantly poor prognosis [75]. Thus, EV miR-335 has been suggested as a novel therapeutic strategy as it was shown to be involved in proliferation and invasion both in vitro and in animal model [76]. Our data suggest an interesting role of EV-based communication that could be explored in the future using our novel 3D co-culture HCC model, which resembles a simplified setting of the TME.

### Secretome profiling in our 3D co-culture supports genes expression profiling and recapitulates *in vivo* signatures

Cellular communication between cancer cells and their surrounding is partially driven by secreted proteins and other soluble factors including cytokines, chemokines, and growth factors. Using antibody microarrays, we determined the levels of different factors and their binding proteins that play a role in insulin pathway, angiogenesis, and cytokine signaling (Fig. 5 B-F). Our results show that secretion of insulin/ IGF pathway proteins IGF-2, IGFBP1, and IGFBP2 is increased by co-culturing HepG2 cells with fibroblasts under 3D co-culture conditions in comparison to mono-cultures (Fig. 5 B & G). IGF-2 is upregulated in several tumors including HCC. Its overexpression was notably detected in HCC patient and was shown to induce liver tumor formation, proliferation and angiogenesis in mice [36]. IGF binding proteins (IGFBPs) are essential in the IGF signaling axis, where they bind with high affinity to IGF-1 and IGF-2, and have been reported in HCC patients [59,77]. Granulocyte-, macrophage-, and granulocyte-macrophage-colony-stimulating factors (G-, M-, GM-, respectively) secreted levels have all increased in 3D co-culture setting in comparison to mono-cultures (Fig. 5 E & G). G-CSF, M-CSF, and GM-CSF have been shown to be involved in liver regeneration, fibrosis, angiogenesis, and initiation and progression of liver cancer [78]. Similarly, levels of factors involved in angiogenic pathway including growth factors, angiopoietins, and matrix metalloproteinases are increased by co-culturing HepG2 cells with fibroblasts under 3D culture conditions (Fig. 5 C & F-H). Despite that the 3D model we are proposing herein is restricted to cancer cells and fibroblasts, present data interestingly shows that our 3D model (group 5 settings) increase the secretion of several cytokines known to orchestrate the cross-talk between the tumor and its immune TME (Fig. 5 D & G-), triggering pro-tumor inflammation and immunosuppression [79]. Our pathway complementation analysis also highlighted two of the enriched genes of group 5 as prognostic markers, namely IGF1R an EGLN3 (Fig. 5 I-J). Both have been reported as indicators of poor HCC prognosis [68,80,81].

In conclusion, we propose a novel 3D tissue culture model of liver cancer that better mimics *in vivo* settings. Compared to the conventional 2D culture, the proposed model exhibits an increase in the expression of genes associated with development, progression, and poor prognosis of HCC. Our results showed that in vivo like conditions cannot be mimicked by simply growing cancer cells as spheroids, but by co-culturing them with 3D fibroblast with which they were able to cross-talk. This was evident by the upregulation of several pathways involved in HCC, and the dramatic jump in secreted factors and surface receptors by co-cultured cancer cells, many of which are also involved in tumor-stroma interactions. We have explored the aspects of HCC our proposed model mimic by combining transcriptome and small scale-secretome analysis, which could be expanded in the future to include proteome analysis. Compared to the conventional 2D culture, the proposed model exhibits an increase in the expression of genes associated with development, progression, and poor prognosis of HCC. Our results correlated with a robust phenotype that better mirrors *in vivo* HCC, from gene expression to prognosis markers, and therefore, a more reliable platform for molecular understanding of HCC and possibly better anti-cancer drug screening platform.

## Acknowledgements

KCG is supported by a CONACYT scholarship (2019-000021-01EXTF-00542). RC is supported by SNF (CRSK-3_190550), BBSRC (BB/N017773/2), and the UZH Research Priority Program (URPP) “Translational Cancer Research”. AA is supported by ZCHS (31R174), UPAR (31S319), and Terry Fox Foundation (21S103).

## SUPPLEMENTARY INFORMATION

**Supplementary data:** materials and methods, and supplementary figures

**Table S1:** Canonical pathways list

**Table S2:** miRNA gene interaction list

**Supplementary document1:** Venn Dr file

**Supplementary document2:** Venn Ur file

