## Supplemental information for "Modelling liver cancer microenvironment: novel 3D culture system as a potential anti-cancer drug screening tool"

**MATERIAL AND METHODS**

***Cell Lines and cell culture***

HepG2 (human Hepatoma cell line) and SV-80 (human fibroblast) were purchased from CLS (CLS GmbH, Germany). HepG2 and SV-80 cells were maintained in high-glucose DMEM medium supplemented with 10% FBS and 1% antibiotic/ antimycotic cocktail (HyClone, UK) at 37°C in 5% CO_2_ humidified incubator. Cells were sub-cultured every 2-4 days using trypsin 0.25%. For 2D mono-cultures, HepG2 and SV-80 cells were cultured at a density of 1X10^6^ in conventional 2D culturing flasks in DMEM media. Flasks were incubated at 37°C in 5% CO_2_ humidified incubator. For generation of HepG2 spheroids, HepG2 cells were trypsinized and resuspended as single cell suspension before being seeded at 1X10^6^ cell density in Corning® Ultra-Low attachment cell culture flasks coated with poly-HEMA (Corning, USA) to facilitate spheroid formation. Plates were incubated at 37°C in 5% CO_2_ humidified incubator and spheroid formation was monitored using inverted microscopy. For generation of 3D culture of fibroblasts, Alvetex strata inserts (Reinnervate, UK) were used. SV-80 cells were trypsinized and resuspended as single cell suspension for cell counting. Alvetex inserts were prepared prior to cell seeding by three washes (1^st^: 70% ethanol, 2^nd^: growth media, 3^rd^: growth media). Inserts were placed in 6-well plate and SV-80 cells were seeded in the Alvetex inserts in a density of 1X10^6^.

***Hypoxia-mimicking conditions***

To mimic the hypoxic microenvironment, 2D cultures of HepG2 cells were cultured as previously described in section 2.2 and treated with 200 and 300 μM of Cobalt (II) Chloride hexahydrate (CoCl_2_) for 6 hours prior to harvesting. The doses and incubation time are based on literature demonstrating HIF1- α maximum induction at 4-6 hours of CoCl_2_ treatment, over a range of doses. Under normoxia, HIF1-α protein is degraded, and hence, serves as a marker for hypoxia. Western blot and immunofluorescence were carried out to assess expression and localization of HIF1-α in CoCl_2_ treated and non-treated cells.

***Morphology Assessment***

To study the morphological differences between 2D and 3D cultures, HepG2 and SV-80 cells were seeded in in 8-well chambers for 2D culture; or in ULA plates and Alvetex inserts, respectively, for 3D culture. 2D cultures of HepG2 and SV-80 were seeded at a density of 2X10^4^ and 1X10^4^ cells/ well in an 8-well chamber, respectively. They were fixed with 100% methanol and stained with crystal violet. 3D culture of HepG2 was seeded at a density of 1X10^6^ and monitored over a period of 5 days. Spheroids were harvested, fixed, and stained with crystal violet and immobilized on agarose pads for imaging. 3D culture of SV-80 was seeded at a density of 0.5X10^6^, then fixed and stained with neutral red. Inserts were unclipped and scaffolds were placed on glass slides for imaging using IX53 inverted microscope (Olympus, Japan).

In addition, immunofluorescence was utilized to better visualize the difference between SV-80 2D and 3D cultures. Cultures were washed with ice-cold 1X PBS, fixed with pre-chilled methanol for 10 minutes at -20°C, permeabilized with 0.5% Triton X-100 PBS, and blocked with 1% BSA solution (in 0.5% Triton X-100 PBS) for 30 minutes at room temperature. Cells were incubated with primary antibody against α-Tubulin (1:100; ab176560) overnight at 4°C, then incubated with Alexa Fluor® 488- conjugated secondary antibody (1:200) for 1 hour at room temperature, and counterstained with DAPI. Antibodies were diluted in 1% BSA. Images were taken using IX53 inverted microscope (Olympus, Japan).

***Cell Viability***

To assess the effects of hypoxia-mimetic agent CoCl_2_ on cellular viability, HepG2 cells were seeded at a density of 5X10^3^ cells/ well in a 96-well plate. Cells were allowed to attach prior to treatment with increasing concentrations of CoCl_2_ (100-400 μM). Cell viability was assessed using CellTiter-Glo Luminescent Assay (Promega, USA), according to manufacturer instructions. Luminescent signals were recorded using GloMax Discover (Promega, USA). The experiment was repeated three times (n=12).

***Western Blotting***

Cells were harvested in ice-cold PBS and then lysed with RIPA buffer containing phosphatase and protease inhibitor and incubated on ice for 30 minutes. Total protein was separated with SDS-PAGE. PVDF membrane was blocked with 5% non-fat dry milk TBST for 1 hour at room temperature. Membranes were incubated with primary antibody against HIF1α (ab1), GAPDH (ab181602), VEGFR1 (ab32152), VEGFR2 (ab134191), TNFR1 (ab68160), TBP (ab220788) for 1 hour at room temperature. All secondary antibodies were diluted in 5% non-fat dry milk TBST. Blots were then visualized using LI-COR C-DiGit Blot Scanner.

***Immunofluorescence***

HepG2 cells were seeded at a density of 1X10^4^ cells/ well in an 8-well chamber and allowed to reach confluency. Cells were incubated with or without CoCl_2_ for 6 hours, after which they were washed with ice-cold PBS and then fixed with cold absolute methanol for 10 minutes at -20 °C. Cells were washed and incubated with 1% BSA blocking solution for 30 minutes at room temperature. Blocking solution was discarded and cells were washed and incubated with antibody against HIF1α (ab1,) at 1:50 dilution at 4 °C overnight. Cells were washed and incubated with secondary antibody prior to counter-staining with DAPI and imaging.

***RNA-Seq Libraries Construction and Sequencing***

For RNA extraction, all groups were prepared as described previously, then collected, washed with 1X PBS, and resuspended in RNAlater stabilization solution before storing at -80°C. Total RNA was isolated from three biological replicates of all groups using RNeasy Mini Kit (Qiagen) following manufacturer’s instructions. Concentration and purity of total RNA was assessed using NanoDrop2000. Quality control of RNA samples was performed with Agilent 2100 Bioanalyzer RNA 6000 Nano Kit, concentrated samples were diluted with RNase-free water prior to bioanalyzer run.

The RNAseq libraries were prepared by DNA Sequencing Center in Brigham Young University. Briefly, KAPA Stranded mRNA-Seq Kit (Kapa Biosystems, USA) was used for capturing poly(A) RNA, converting it to cDNA, A-tailing, and Adapter ligation. Fragments carrying appropriate adapter sequences were amplified to yield mRNA-Seq libraries. KAPA Library Quantification Kit (Kapa Biosystems, USA) was used for libraries quantification prior to Illumina sequencing using high-throughput Illumina HiSeq sequencing system (Illumina, USA).

***Alignment and Analysis of Illumina Reads***

Briefly, reads obtained from Illumina were aligned to Homo sapiens GRCh38.p2 reference genome using tophat2 v2.1.0. Following alignment and annotation, read counts were generated using HTseq count. Read counts were used to generate principle component analysis (PCA) plot and hierarchical cluster heatmaps, using the web-based tool ClustVis (<https://biit.cs.ut.ee/clustvis_large/>), for clustering of multivariate data. Triplicates of each group are collapsed by taking the mean, and rows were scaled using vector scaling method. Due to limitation on input data size, 2400 genes were selected randomly for generation of heatmap in Figure 2.

***Differential Gene Expression Analysis***

RNA-seq 2G (<http://54.243.174.165:3838/rnaseq2g/>) was used to perform analysis of differential gene expression using read counts between group 1 (2D HepG2 under normoxia), group 2 (2D HepG2 treated with CoCl_2_), group 3 (2D co-culture HepG2 treated with CoCl_2_), group 4 (3D HepG2), and group 5 (3D co-culture HepG2). Counts were normalized using DEseq method, and differential expression was determined using DEseq2 method. Obtained DEGs with false discovery rate (FDR) <0.05 and exhibiting a fold change ≤ -2and ≥ 2 were identified as significant DEGs.

***Quantitative polymerase chain reaction***

For the purpose of validating genes of interest, quantitative polymerase chain reaction (qPCR) was carried out on samples from groups 1 and 5. Total RNA was converted to cDNA using GoScript™ Reverse Transcription System, according to manufacturer's instruction (Promega, USA). GoTaq® qPCR Master Mix (Promega) was used to perform qPCR on QuantStudio 5 Real-Time PCR System (Applied Biosystems) using gene-specific primers purchased from Macrogen (Macrogen Inc.). 18S rRNA was used for data normalization due to its unchanged expression in groups 1 and 5. Comparative C_T_ method (2^-ΔΔC^_T_) was used to determine fold change in expression of target genes between group 5 and group 1, according to the following equation: 2^-ΔΔC^_T_ =[(C_T_ gene of interest - C_T_ internal control) group 5 – (C_T_ gene of interest - C_T_ internal control) group 1]. The cycling parameters recommended in GoTaq® qPCR Master Mix manual were used. Briefly, Hot-Start Activation was carried at 95°C for 2 minutes, 40 cycles of denaturation at 95°C for 15 seconds, followed by annealing/ extension at 60°C for 60 seconds, and then dissociation at 60–95°C.

***Gene Set and Gene Ontology Enrichment Analyses***

Gene set enrichment analysis (GSEA) was carried on sets of upregulated and downregulated genes using canonical pathways ontology on eXploring Genomic Relations (XGR) web version (http://galahad.well.ox.ac.uk:3030/). Enriched terms were tested for significance using the Hypergeometric test, and only terms with FDR < 0.05 were considered. Gene ontology (GO) enrichment analysis was carried out using Biological Networks Gene Ontology (BiNGO) App [28] in Cytoscape, an open source software platform for network data integration, analysis, and visualization [29]. Based on Hypergeometric significance test, corrected multiple testing using Benjamini and Hochberg FDR < 0.05, and using biological process ontology, relevant enriched terms were identified by BiNGO. InteractiVenn (<http://www.interactivenn.net/>) was used to generate a Venn diagram of relevant GO terms by uploading sets of their associated genes [30].

***Networks Analysis***

The web platform Network Analyst (<http://www.networkanalyst.ca/>) was used to generate global gene expression PCA plot. ORA Heatmaps were done based on the top enriched pathways using KEGG database.

***PCA plot analysis***

Principal component analysis (PCA) was performed in Multibase 2015 - Excel add in (Numerical Dynamics, Japan, 2015). Here, the overall shared similarities and differences of the group comparisons are analyzed and visualized in a plot. As an input, the different group comparisons were merged in a matrix based only on the differentially expressed molecules (includes ID and LFC) shared across all the group comparisons (N=350).

***Functional Analysis***

Functional analyses were performed on differential expressed genes with FDR <0.05 from the different group comparisons (G2 vs G1, G3 vs G1, G4 vs G1 and G5 vs G1). Gene onthology (GO) and pathway clustering analysis were performed using the ClueGO, Cytoscape plug-in; pathway analyses were based on the Kyoto encyclopedia of genes and genomes (KEGG) and WikiPathways annotations built-in ClueGO [31]. CluePedia Cytoscape plug-in, was used on ClueGO outcome to show the molecules tighten to the highlighted nodes/networks [32]. ClueGO/CluePedia analyses followed default parameters (or otherwise in-text specified). Term convergence among GO, KEGG and Wikipathways terms assures the validity of the model created to each of the group comparisons.

***Pathway analysis***

Pathway visualization and analysis were performed in PathVisio 3.3.0. These analyses were based on the Wikipathways human collection. Primary analyses were carried out in order to visualize how the most significant molecules included in ClueGO/CluePedia analysis fit on the most significant pathways identified through this functional clustering. To import this data into Pathvisio, as an input the following data were included per molecule in a table format: EnsEMBL ID, FDR and logFC. After pathway mapping and identification of these molecules within the pathway maps of interest, all the molecules with an FDR below 0.05 were included into Pathvisio in order to fill the gaps within the focused pathways and proceed to identification of trends in regulation. LogFC values from 1 to -1 are represented and visualized in a color gradient from red to green respectively.

***Growth Factor Antibody Array***

Secreted growth factors, angiogenesis factors, and cytokines in conditioned media of 3D mono- or co-cultures were analyzed using Human Growth Factor Antibody Array kit (ab134002, abcam), human Angiogenesis Antibody Array kits (ab169808/ ab134000, abcam). Briefly, membranes were incubated with the blocking solution for 30 minutes at room temperature. After which membranes were washed and incubated with 2 mL of conditioned media collected from indicated groups overnight at 4 °C. Membranes were washed and incubated with Biotin-Conjugated Anti-Cytokines overnight at 4 °C. The membranes were the incubated with 2 mL of HRP-Conjugated Streptavidin for 2 hours at room temperature. Membranes were washed prior to incubation with detection buffer for 2 minutes and imaged using ChemiDoc imaging system (Biorad). Mean pixel intensity were determined using ImageJ.

**Supplementary Figures**

**Fig. S1.** Uncropped blots of Fig. 1

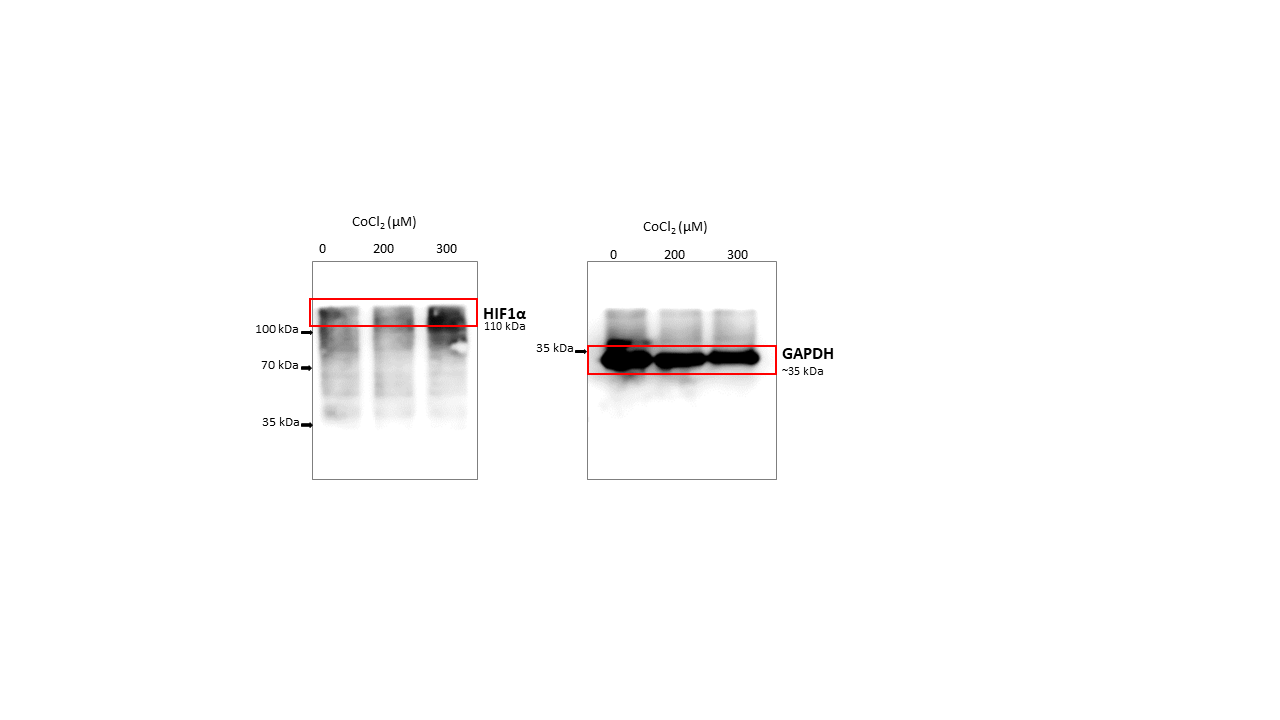

**Fig S2.** a) Schematic design of experimental outline. b) levels of different secreted factors in 3D mono- and co-cultures analyzed by antibody arrays.

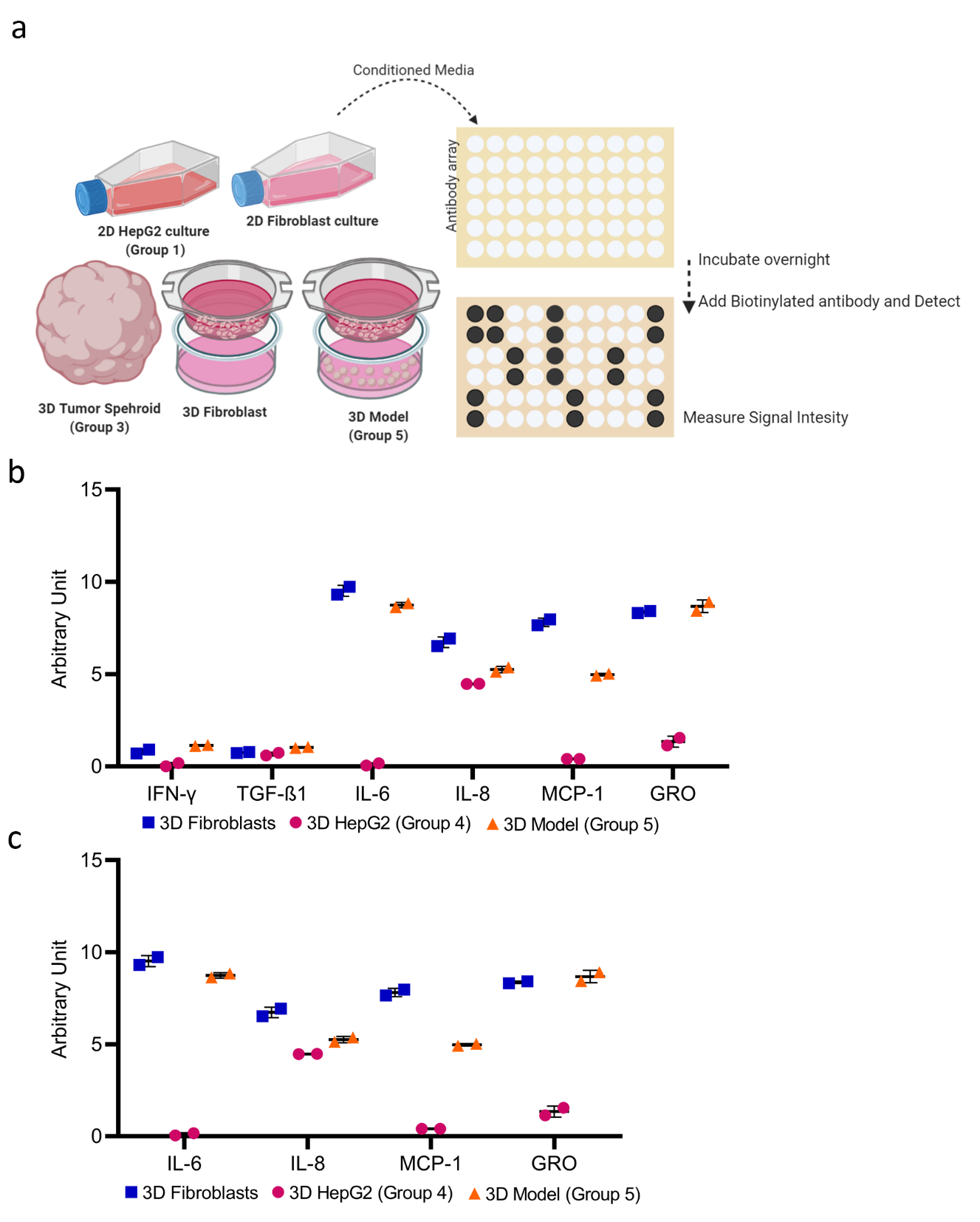

**Figure S3.** Uncropped blots of Fig. 5

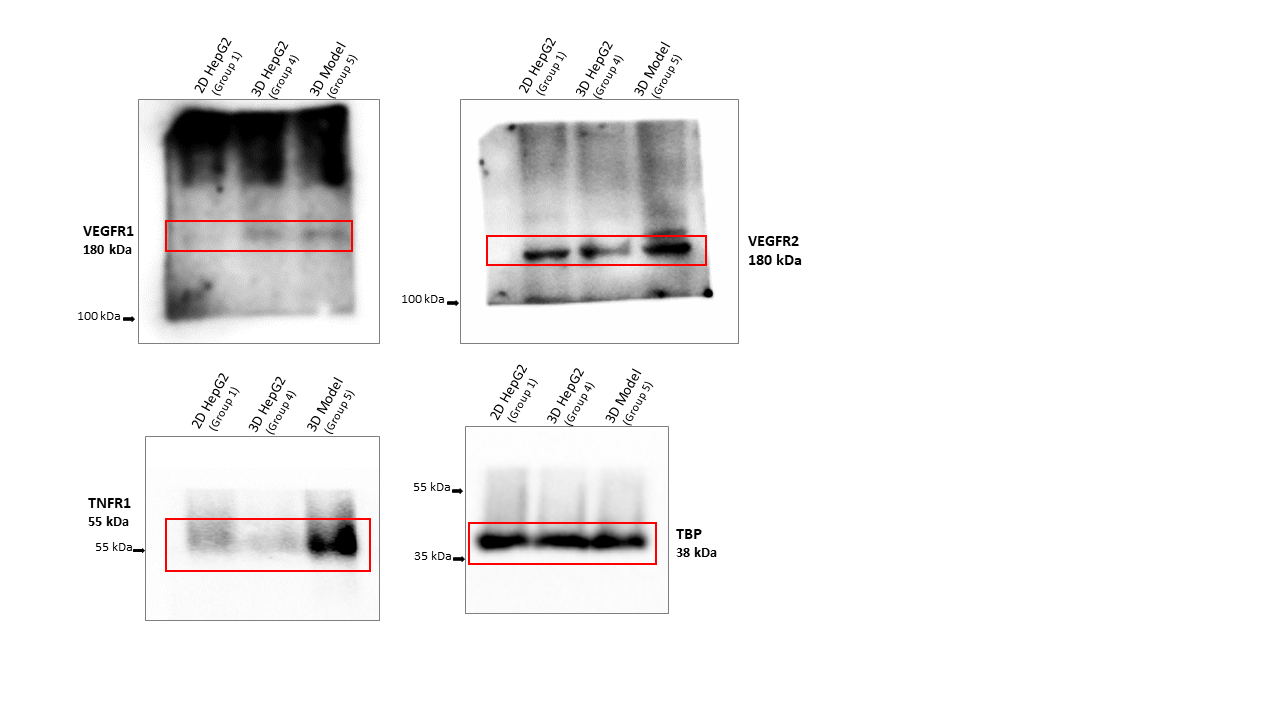

**Figure S4.** **Pathway complementation analysis highlights regulatory miRNAs.**
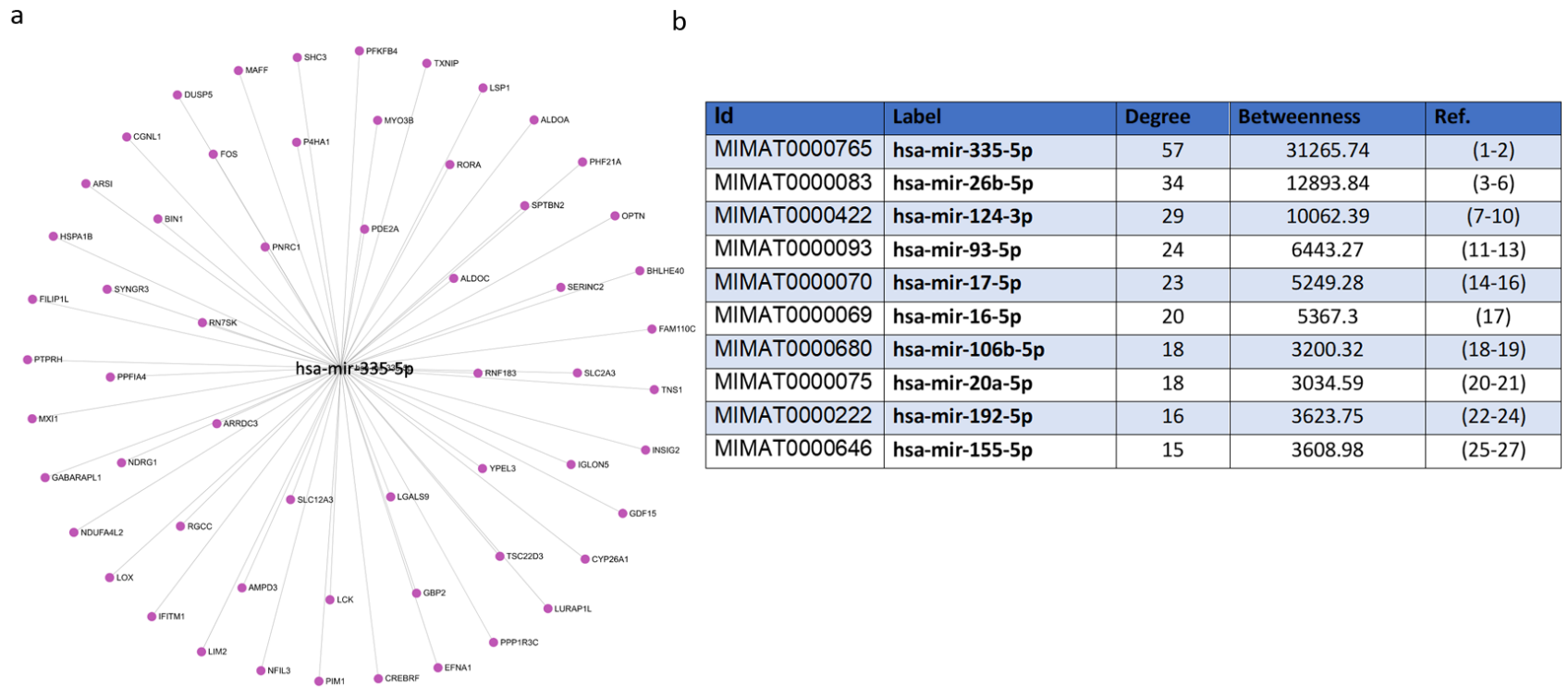

a) miR-335 and its gene interactions. b) table of top 10 miRNAs in the miRNA-gene network of group 5 and literature references [1–27].
References
